# Different timescales of neural activities introduce different representations of task-relevant information

**DOI:** 10.1101/2024.07.23.604720

**Authors:** Tomoki Kurikawa

## Abstract

Recent findings indicate significant variations in neuronal activity timescales across and within cortical areas, yet their impact on cognitive processing remains inadequately understood. This study explores the role of neurons with different timescales in information processing within the neural system, particularly during the execution of context-dependent working memory tasks. Especially, we hypothesized that neurons with varying timescales contribute distinctively to task performance by forming diverse representations of task-relevant information. To test this, the model was trained to perform a context-dependent working memory task with a machine-learning technique. Results revealed that slow timescale neurons maintained stable representations of contextual information throughout the trial, whereas fast timescale neurons responded transiently to immediate stimuli. This differentiation in neuronal function suggests a fundamental role for timescale diversity in supporting the neural system’s ability to integrate and process information dynamically. Our findings contribute to understanding how neural timescale diversity underpins cognitive flexibility and task-specific information processing, highlighting implications for both theoretical neuroscience and practical applications in designing artificial neural networks.

## 1 Introduction

The neural system is a complex network composed of components operating at various timescales. Slow synaptic dynamics, also known as synaptic plasticity, establish elaborate synaptic connectivity among neurons, while fast neural dynamics are responsible for performing cognitive tasks. Recent experimental studies have revealed a significant diversity in timescales of neuronal activities across different cortical areas[19, 15, 14] and within individual cortical areas[20, 4, 16, 2]. Despite the widespread observation of these multiple timescales in neural activities, the impact of neurons with different timescales on information processing within the neural system remains unclear.

Several modeling studies have demonstrated that neural networks with neurons operating at multiple timescales can generate compositional sequences[21, 11, 13]. In these models, slow neural activity acts as a “glue” that binds neural patterns formed by fast neural activities to create complex sequential activities. However, the role of these multiple timescales in performing complex cognitive tasks, such as context-dependent working memory tasks, beyond sequence generation, is not well understood.

In the present study, we focus on the different representations of task-relevant information by neurons with varying timescales during the performance of a context-dependent working memory task. Previous experimental studies[18, 17] have shown distinct neural representations corresponding to context signals, sample cues, and go cues in context-dependent tasks. These diverse representations are crucial for integrating information to perform context-dependent tasks. We hypothesize that neurons with different timescales are fundamental to the emergence of these varied representations.

To test this hypothesis, we constructed a recurrent neural network model composed of neurons with multiple timescales and trained it to perform a context-dependent working memory task. We found that slow timescale neurons represent the current context and maintain this representation throughout the trial, whereas fast timescale neurons instantaneously represent the applied stimuli.

## 2 Task and Model

### 2.1 Network model

To investigate how the dynamics with different time scales of neural activities perform a cognitive task, we build a recurrent neural network model trained to perform a context-dependent working memory task (Fig. 1). The recurrent network has *N* rate coding neurons following the equation

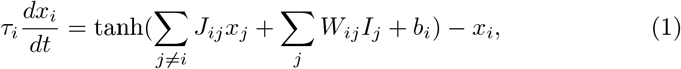

where *x*_*i*_ (*i* = 1,,, *N*) is the activity of *i*-th neuron with its time scale *τ*_*i*_ and *J* is a connectivity matrix in the recurrent network. *W* is a connectivity matrix from the input neurons to the recurrent network and *I* represents the activities of input neurons, which will be explained later. Now we have fast and slow neurons: the timescales of the neurons *τ* are set to

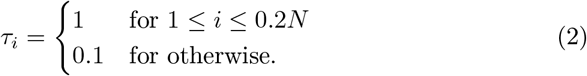

*J* is an all-to-all connectivity that is modified through learning. The connectivity structure does not depend on different timescale neurons.

**Fig. 1.**
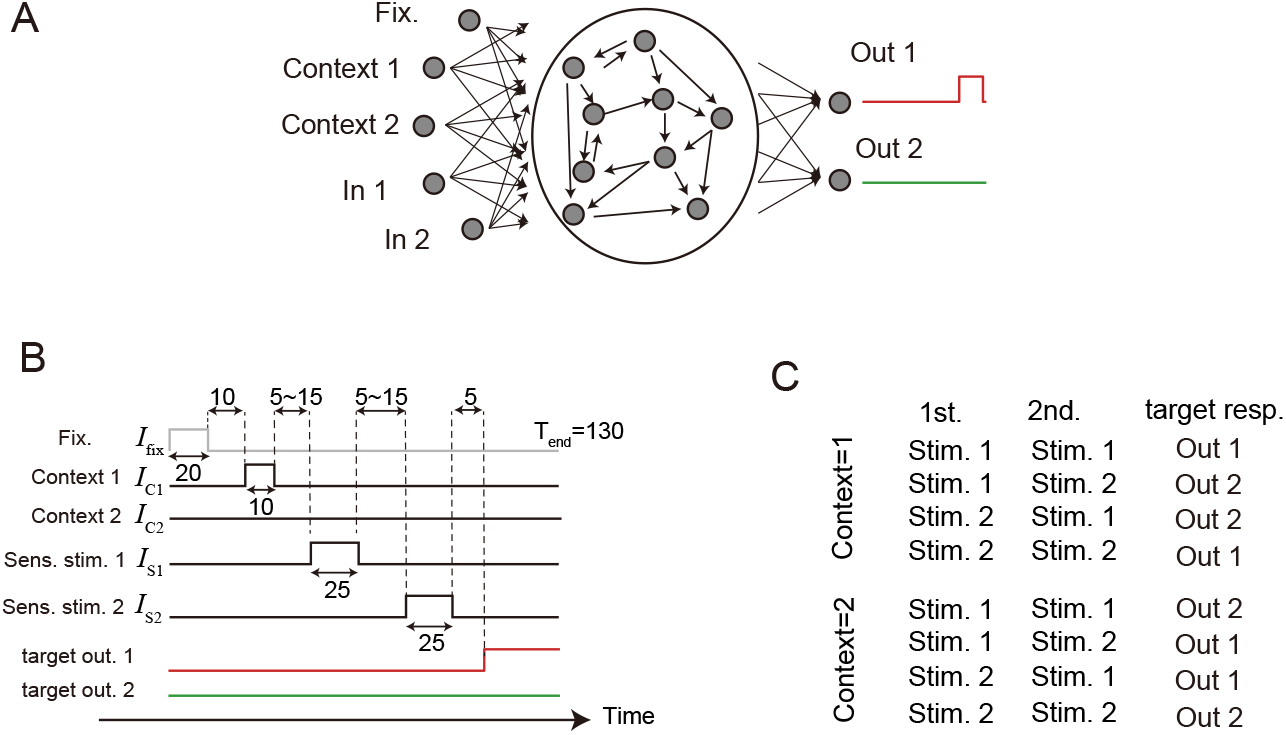
Schematic image of our model and task. A: Network image. B: Task procedure showing a trial with context 1, 1st stimulus being stimulus 1, and 2nd stimulus being stimulus 2. C: A table showing the mapping between the inputs and target outputs, depending on the context.

The recurrent network receives five input neurons representing fixation cue (fix. neuron), context cues (two context neurons), and sensory stimuli (two stimulus neurons). The temporal patterns of activation of these input neurons are given according to a task procedure explained later. These input neurons are connected to the neurons in the recurrent network through the connectivity *W* which is also modulated through learning. The activities of the recurrent neural network are read-out by two output neurons that correspond to the behavior of the network.

### 2.2 Task

The task the network has to perform is a context-dependent working memory task as shown in Figs. 1B and C. In the task, five input neurons are activated sequentially. First, the fix. neuron is activated, followed by the activation of one of the two context neurons. Then, the sensory stimulus neurons are activated twice sequentially with delayed time in four different temporal patterns: stim 1 - stim 1, stim 1 - stim 2, stim 2 - stim 1, and stim 2 - stim 2. Depending on the context given by activation of either context neuron, the network has to respond differently to these sensory stimulus patterns as shown in Fig. 1C. Under context 1 (context neuron 1 is activated), the output neuron 1 is supposed to activate when the same stimulus is applied twice, otherwise, the output neuron 2 should be activated. Conversely, in the context 2, the required association between sensory stimulus patterns and output patterns is the opposite.

The durations of the activations of neurons and their intervals are shown in Fig. 1B. When the input neuron is activated, its activity (for instance *I*_fix_ for the fix, neuron) is set at 1. The input to the recurrent network, *I* in Eq. 1, is composed of activities of all input neurons *I* = (*I*_fix_*I*_C1_*I*_C2_*I*_S1_*I*_S2_)^*T*^.

The networks are trained so that the targeted output neuron activates 5 unit-times after the second sensory stimulus is disactivated. The activities of the output neurons 1 and 2 are denoted by *out*_1_ and *out*_2_, respectively, and are given by

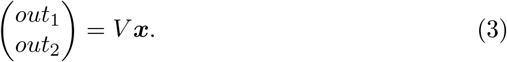

Here, *V* is an output connectivity from the recurrent network to the output neurons which is plastic through learning.

### 2.3 Training method

We use the standard backpropagation through time with Adam optimizer in pytorch to train the network. All connectivity *W, J, V* and the bias are modulated through learning. The target patterns are shown in Fig. 1B. The activities of output neurons are required to be zero before Go period that starts 5 unit-time after the offset of the 2nd sensory stimulus. Then, the target output neuron is required to be one, whereas the non-target output neuron is required to be still zero. The Loss function is the mean square error between the target output behaviors and actually generated output behaviors. The parameters used in this training are shown in Table.

**Table 1.** Table: parameters in the model.

| parameter | value |
| --- | --- |
| the number of neurons in the recurrent network, $N$ | 100 |
| learning rate in Adam | 0.001 |
| coefficient of inertia term in momentum update | 0.9 |
| coefficient of inertia term in RMSprop update | 0.999 |

## 3 Results

### 3.1 Neural dynamics after learning

First, we briefly examine whether the learning is actually complete. Figure 2 shows the learning curves across ten realizations of networks with different initial values in their connectivity matrices. All of the learning curves decrease and eventually reach almost zero, indicating that the networks have completed the context-dependent working memory task.

**Fig. 2.**
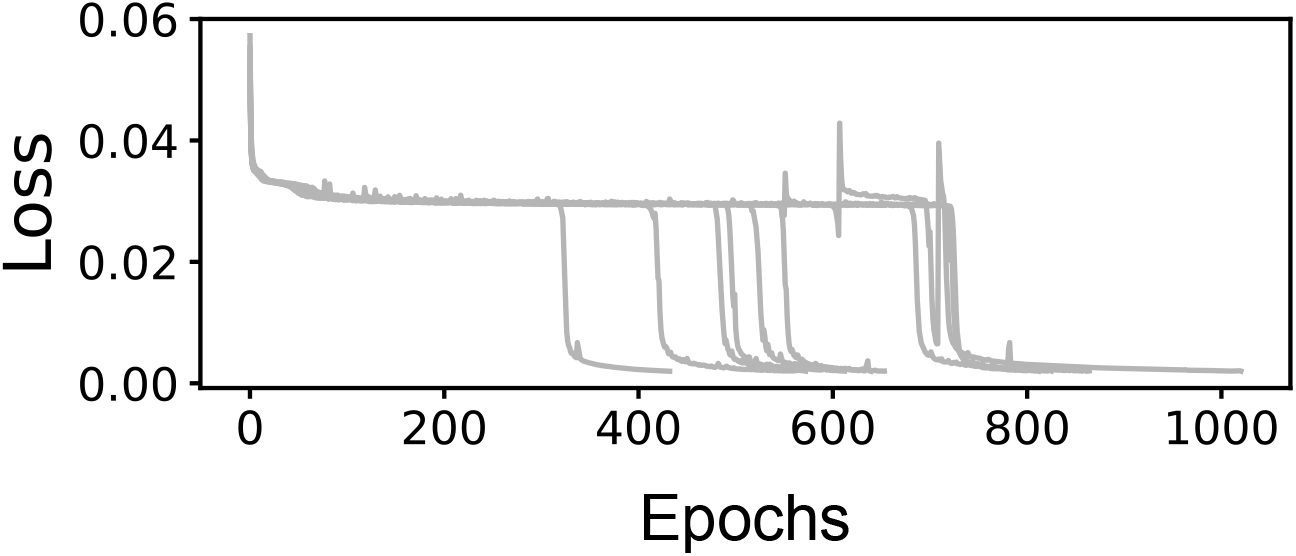
Learning curves. Learning processes for ten different realizations of initial connectivities.

Next, we analyze the neural dynamics after training is completed. Figure 3 exhibits the activities of the neurons in the recurrent network and the output neurons for different sensory input patterns under context 1. In this task, the output neurons are required to respond to the patterns of two sucessive sensory inputs, not to the last input itself. In fact, the output neurons in the trained network successfully generate appropriate patterns.

**Fig. 3.**
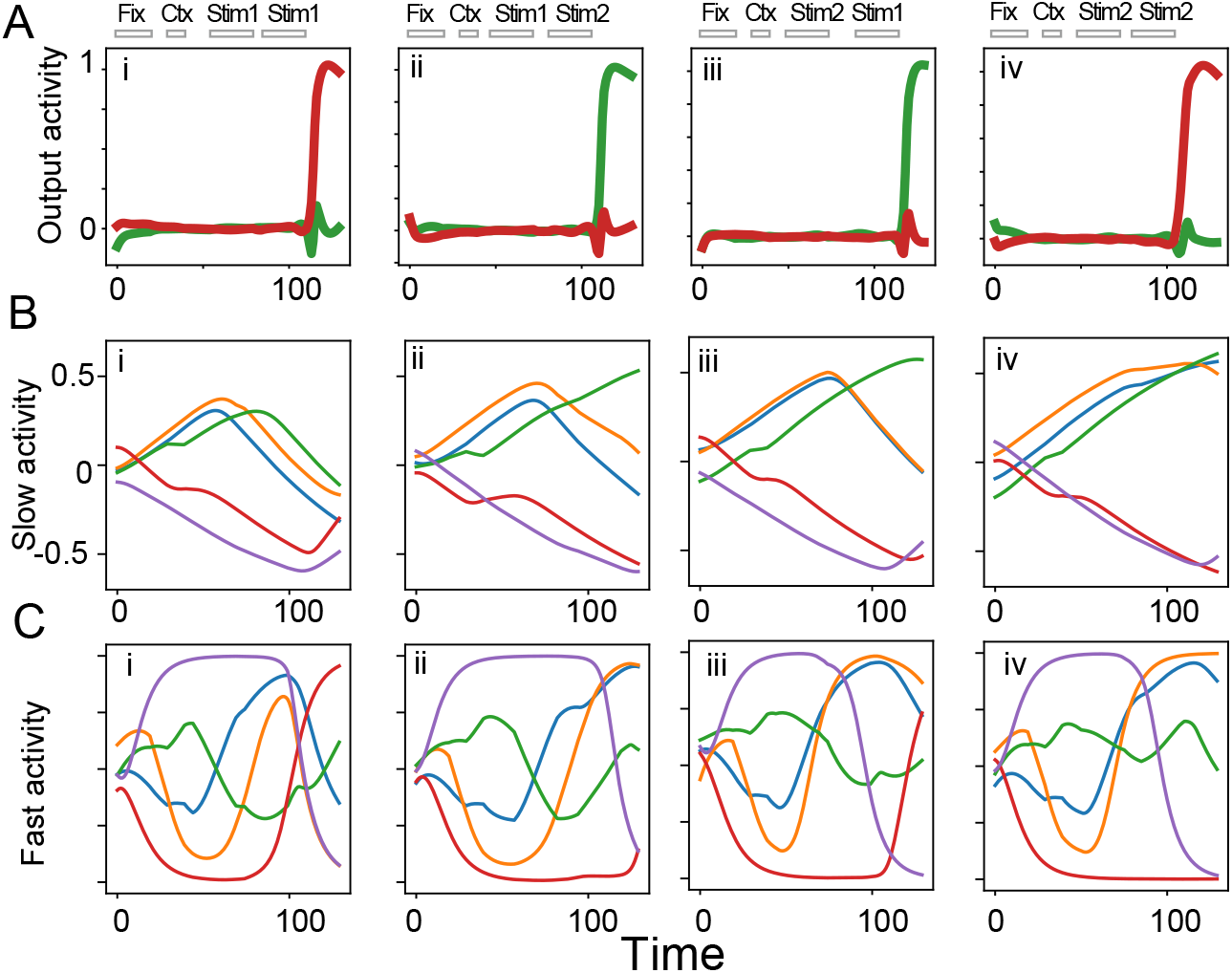
The neural activities in the context 1 after training. A: The activities of two output neurons are plotted for different four input patterns. The application time of all the external cues is exhibited by bars at the top of the panels. B: The activities of the slow neurons are plotted. The activities of different neurons are represented by different colors. C: The activities of the fast neurons are plotted in the same manner as in B.

### 3.2 Trajectories of neural population

To analyze the neural dynamics at the population level, we visualized the neural trajectories by projecting on the three largest principal components (PCs). We used all neurons in the recurrent network for the analysis. Figure 4A illustrates the neural trajectories as the network performs the task under context 1. The neural states evolve from the initial states to the states with the context 1 cue applied (magenta circle), the states with the first sensory stimuli, and the states with the second stimuli applied (states with the fixation cue are not shown for clarity). Finally, the neural states reach the endpoints of the time series that correspond to decision points. In the two trials represented by the blue and green lines, where the first and second stimuli are the same, the network gives the activation of output 1. Thus, the network successfully completes the task. However, these trajectories do not appear to converge to the same state in this PC space because the states in the recurrent neural network are different even in the same output trails.

**Fig. 4.**
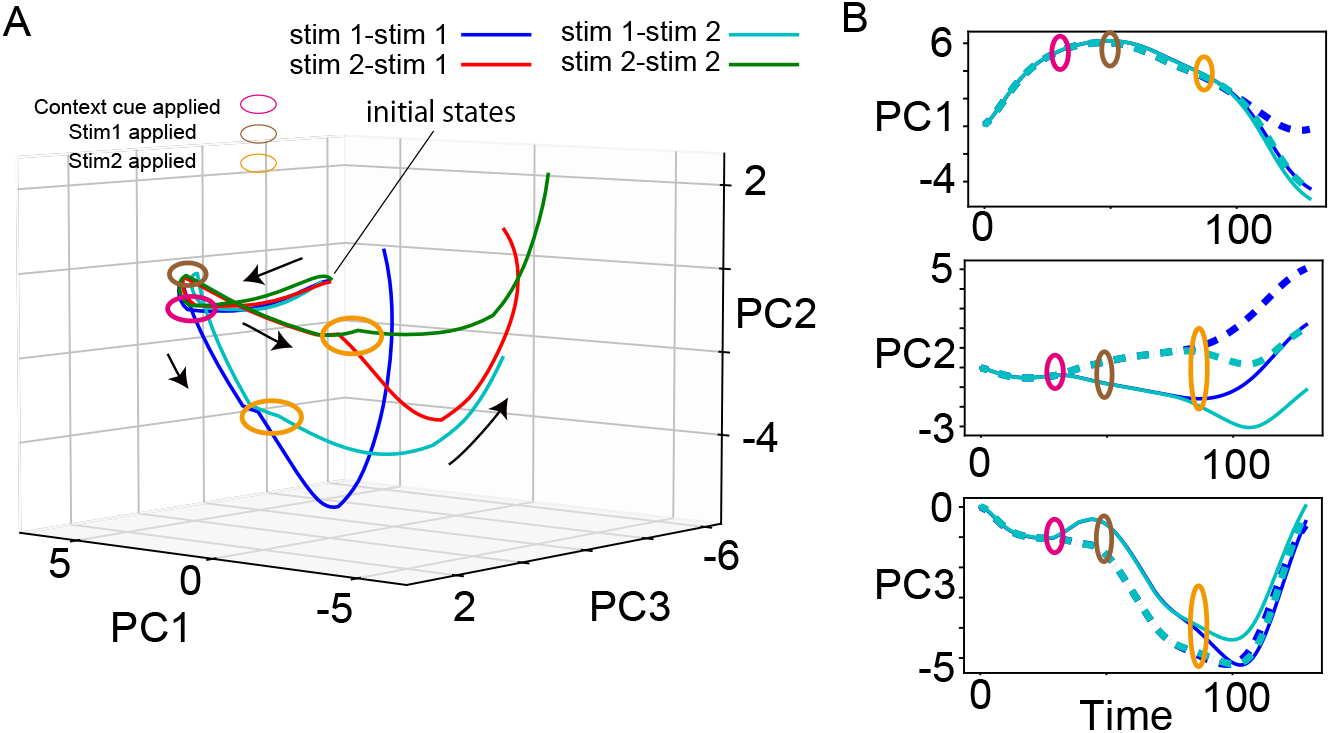
Neural dynamics projected into PC space. A: The neural dynamics for context 1 are plotted into the largest three PCs for four patterns of the applied inputs in different color lines as shown above the panel. The states marked by the colored circles correspond to the states with the application of the cues. B: The neural dynamics are projected into the largest three PCs separately, for two different input patterns under contexts 1 and 2 in the same way as in A. The neural activities under context 1 are illustrated in solid lines, while those under context 2 are in dotted lines.

So far, we have analyzed how the neural trajectories evolve under context 1. How is the context represented in the neural trajectories? To answer this question, we plot four neural trajectories for the two sensory stimulus patterns, stim. 1 - stim.1 and stim.1 - stim. 2, under context 1 and context 2 in Fig. 4B, by projecting them separately onto the three largest PCs. The neural trajectories projected on the largest PC show almost no difference between conditions and between contexts as well as those on the 3rd PC. In contrast, the trajectories projected on the second-largest PC clearly differ between context 1 and context 2. The difference emerges at the onset of the context cue and enlarges until the end of the trials. Thus, the 2nd PC reflects the information about the current context. In this way, the population of neurons encodes the context information.

Next, we ask whether there are different representations between fast and slow neurons. To clarify this possible difference, we analyzed the neural dynamics composed by the fast neurons and those by the slow neurons, separately. Figure 5A exhibits the dynamics of the slow neurons projected onto the three largest PCs. The dynamics projected onto the largest and 3rd largest PCs differ dramatically depending on the context conditions after the onset of the context cue. In contrast, the dynamics of the fast neurons demonstrate the relatively small difference between the contexts 1 and 2 as shown in Fig. 5B: The largest PC dynamics show no difference and the second and third PC dynamics show a small difference between contexts 1 and 2.

**Fig. 5.**
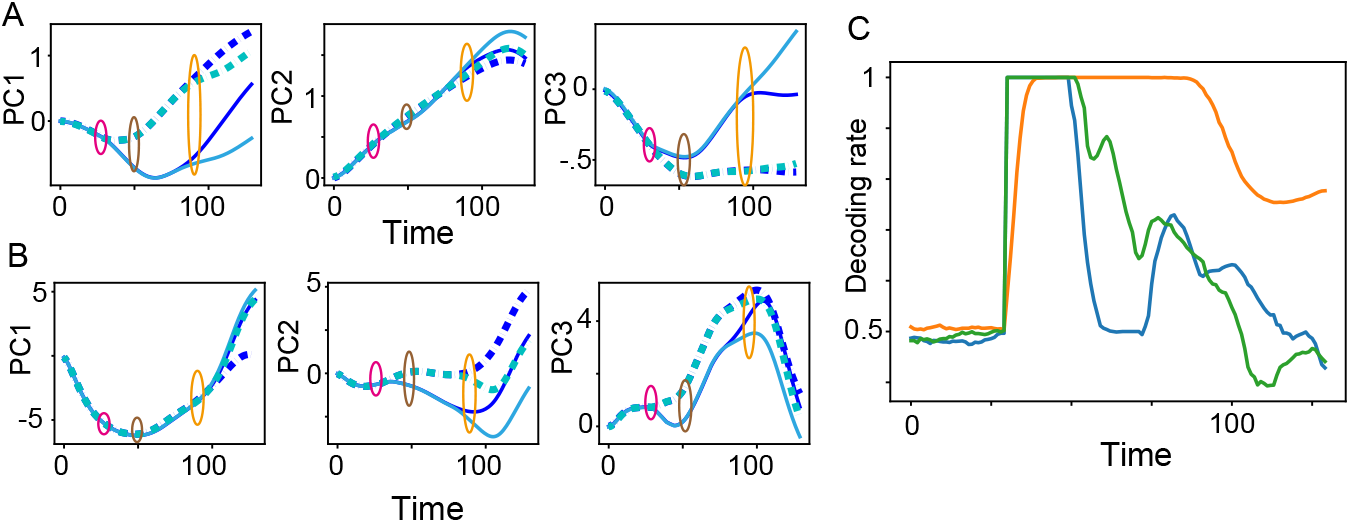
The slow and fast neural dynamics encode the context information differentially. A and B: The neural activities for four different conditions are shown in the same manner as in Fig. 4. C: The decoding rate of the context information in different neural populations. The green, blue, and orange curves represent the rate calculated by all, fast, and slow neurons, respectively.

To quantify the difference in the representations of the contexts between the fast neurons and the slow ones, we used linear discrimination analysis (LDA) and computed the decoding rate of which of the contexts was applied. LDA identifies the axis *w*_*LDA*_ that maximizes the difference between the mean values of two groups of high-dimensional states projected on this axis normalized by the variance of each group projected on this axis: 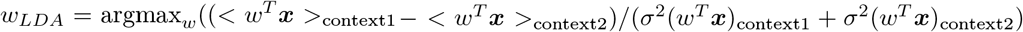. Here, *w*^*T*^ is a transpose of the vector *w* and ***x*** is a vector of the neural state. < … >_context1_ is the mean value across the ensemble under context 1, while *σ*^2^(…)_context1_ is a variance across the ensemble under context 1. We computed *w*_*LDA*_ by using the neural states at the end of the context input application. For this *w*_*LDA*_, we examined to decode which context is applied from the neural state. When the neural state *x* satisfies *w*^*T*^ ***x*** ≤ (< *w*^*T*^ ***x*** >_context1_ + < *w*^*T*^ ***x*** >_context2_)*/*2 for < *w*^*T*^ ***x*** >_context1_ ≤ < *w*^*T*^ ***x*** >_context2_, we infer this state under the context 1. We calculated the success rate of decoding by using three populations of neurons at each time: all neurons, the fast neurons, and the slow neurons. Figure 5C exhibits the success rate of decoding for these three types of neural dynamics. The decoding rates for all neurons and the fast neurons increase temporarily to 1 during only the application duration of the context stimulus. Then, the rate dropped rapidly. On the contrary, the rate for the slow neurons takes the high value persistently after the offset of the context stimulus. Thus, the slow neurons keep the context information after the application of the context input and contribute to performing this context-dependent working memory task.

### 3.3 Cell level representations: slow neurons represent context

Finally, we explore how the context and other information are encoded at single-cell level. To quantify how much information is encoded in each neuron, we computed a selectivity index for a neuron *i* regarding input cue *α* as

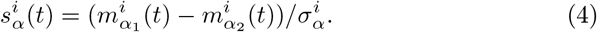

*α* represents the context input, the 1st sensory stimulus, or the 2nd one. For *α* =context input (for *α* = the 1st or 2nd sensory stimuli), *α*_1_ and *α*_2_ specifically represent context 1 (sensory stim. 1) and context 2 (sensory stim. 2), respectively. 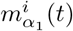 is a mean value of the activities of the neuron *i* at time t across the trials in *α*_1_ condition. 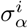 means the standard deviation of the activities of the neuron *i* across all trials.

Calculation of the selectivity index allows us to classify neurons into groups depending on the information they encode. Figure 6 illustrates some representative examples of the selectivity index of different groups of neurons. A group shows the transient information encoding, as shown in Fig.6A(i) and B(i). The selectivity index of this group increases and decreases transiently at the time of the application of the corresponding input. First, the selectivity index of the context information is transiently activated, that of the 1st stimulus information, and, finally that of the 2nd stimulus is activated. In contrast to the expectation, the transient-encoding group of neurons includes the slow neurons, not only the fast neurons.

**Fig. 6.**
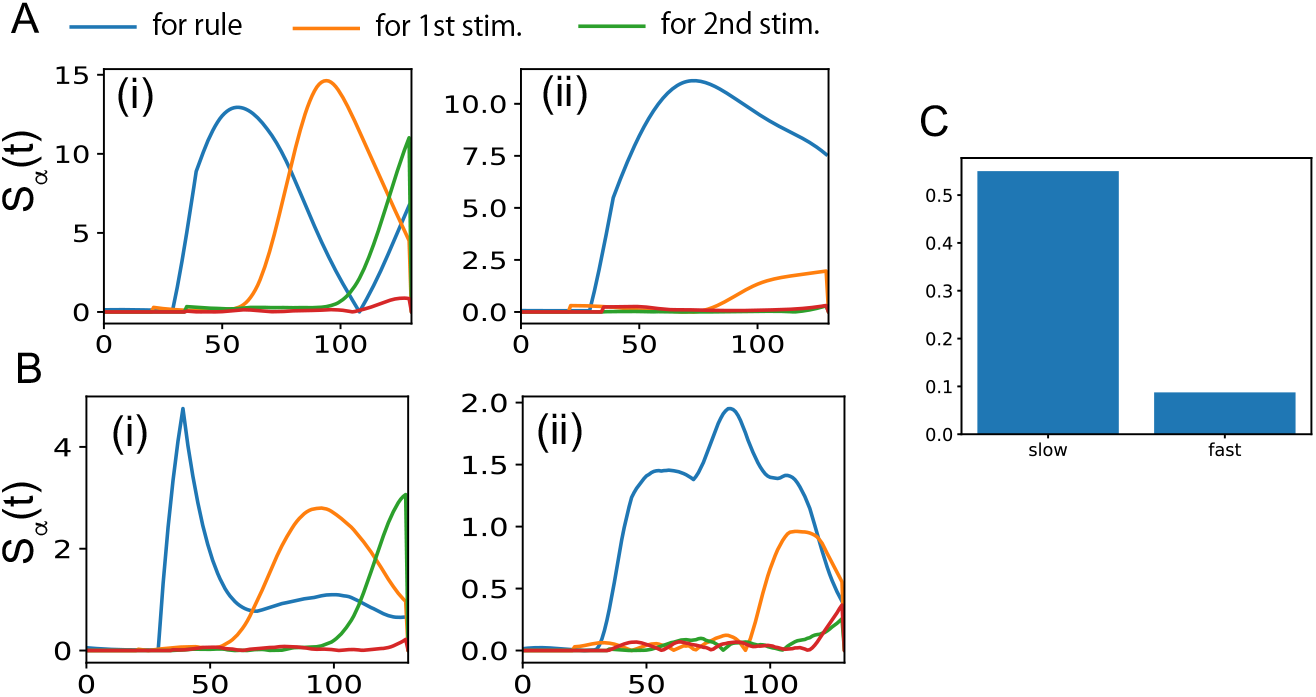
The selectivity index of each neuron. A and B Time series of the selectivity index for the context, the 1st stimulus, and the 2nd stimulus are shown for four different neurons. 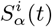 is represented for *α* = context, 1st stim. and 2nd stim. in blue, orange, and green, respectively. 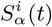 is obtained from the slow neurons in A, while those from the fast neurons in B. Panels AB(i) demonstrate the transient encoding neurons and AB(ii) do the persistent encoding neurons. C The ratio of the context-encoding neurons in the slow and fast neuron populations are plotted.

The other group of neurons shows persistent information encoding, as shown in Fig.6A(ii) and B(ii). In this group, the selectivity index for the context information rises at the onset of the context stimulus and remains the high value at the end of the trial. The selectivity index for other information also increases, but the increase is much smaller than that of the context information. This group includes not only the slow neurons, but also the fast neurons.

We counted the number of persistent encoding neurons for the context stimulus, because such neurons are expected to contribute to the context coding shown in Fig. 5A(ii) and B(ii). We defined the context encoding neuron as a neuron showing that its selectivity index for the context is highest from the beginning to the end of the trial. The number of these neurons is computed and plotted for the fast and slow neurons, separately, in Fig. 6C. More of the slow neurons show the persistent encoding of the context than the fast neurons. This result means that, at the single-cell level, there are more neurons with slow timescales showing context-selective activities.

## 4 Discussion

We demonstrated that the slow and fast neurons differentially represented the task-relevant information. More the slow neurons exhibited the context-selective activities that persisted throughout the trial. In contrast, the fast neurons showed transient activities corresponding to different signals.

Diversity of the timescales of neural activities is widely observed in single areas, such as lateral intraparietal cortex[20], dorsolateral prefrontal cortex[4], orbitofrontal cortex, and anterior cingulate cortex[2]. In these studies, the timescales were estimated by the auto-correlation of the neural activities in the resting state and the neurons with different timescales showed the different behaviors For instance, the studies[20, 3] slowed the slow timescale neurons maintained their activities during the delay period in the working memory task, whereas the fast neurons did not and showed an instantaneous response to the cues. Our study gives a potential mechanism underlying the emergence of such different representations in different timescale neurons.

Diverse representations of multiple task-relevant signals are necessary to perform complex tasks. Actually, besides the multiple timescales, the several characteristics of neurons are related to different representations: hippocampal neurons projecting to different cortical areas encoded different information, such as anxiety and goal-related excitation[6]. Different types of inhibitory neurons in the medial prefrontal cortex demonstrated distinctive roles in the working memory task[9]. Integration of these characteristics not implemented in our current model with different timescales might step ahead toward divisions of single neurons.

The neural systems with slow-fast neurons have been analyzed in the generation of the sequential neural patterns[21, 13, 12, 11, 10] and accurate Bayesian inference[7]. Introducing the slow neurons allows the neural network to produce complex sequences composed of fast neural activity patterns such as a history-dependent sequence such as ABABC in which a pattern C should appear only after twice of AB. However, these mechanisms are not available to perform cognitive tasks such as the context-dependent working memory task (but see [12, 11, 10]). Further, the connectivity across the slow and fast neurons is well-designed to generate sequential patterns. In contrast to these studies, our present study shows that differentiation of representations across neurons emerges through training the cognitive tasks without specific network motifs.

Finally, we discuss the limitations of our model. In our model, two timescales of neural activities are explicitly implemented. whereas, some studies[5, 1, 8], suggested that the diversity of the timescales emerges depending on the connectivity [5, 1] and through learning[8]. The interaction between the emergence of the multiple timescales and their diverse representations is quite an important point for understanding multiple timescales neural systems, which is beyond our present paper. However, our study gives an base model to build a model with change in the timescale of neural activities.

To sum up, the present study demonstrated that, in the network with slow and fast neurons, the different types of neurons show different representations in the context-dependent tasks. The slow neurons prefer to represent the context information that persists during the entire trial so that the context-dependent neural dynamics are generated even in the same sensory stimuli.

## Acknowledgments

This study is funded by the special grant of Future University Hakodate.

## References

1. Bernacchia, A., Seo, H., Lee, D., Wang, X.J.J.: A reservoir of time constants for memory traces in cortical neurons. Nature Neuroscience 14(3), 366–372 (mar 2011)

2. Cavanagh, S.E., Hunt, L.T., Kennerley, S.W.: A Diversity of Intrinsic Timescales Underlie Neural Computations. Frontiers in Neural Circuits 14(December), 1–18 (2020)

3. Cavanagh, S.E., Towers, J.P., Wallis, J.D., Hunt, L.T., Kennerley, S.W.: Reconciling persistent and dynamic hypotheses of working memory coding in prefrontal cortex. Nature Communications 9(1) (2018)

4. Cavanagh, S.E., Wallis, J.D., Kennerley, S.W., Hunt, L.T.: Autocorrelation structure at rest predicts value correlates of single neurons during reward-guided choice. eLife 5(OCTOBER2016), 1–17 (2016)

5. Chaudhuri, R., Bernacchia, A., Wang, X.J.: A diversity of localized timescales in network activity. eLife 3, e01239 (2014)

6. Ciocchi, S., Passecker, J., Mikus, N., Klausberger, T., Malagon-Vina, H., Mikus, N., Klausberger, T.: Selective information routing by ventral hippocampal CA1 projection neurons. Science 348(6234), 560–563 (may 2015)

7. Ichikawa, K., Kaneko, K.: Bayesian inference is facilitated by modular neural networks with different time scales. PLOS Computational Biology 20(3), e1011897 (mar 2024)

8. Khajehabdollahi, S., Zeraati, R., Giannakakis, E., Schäfer, T.J., Martius, G., Levina, A.: Emergent mechanisms for long timescales depend on training curriculum and affect performance in memory tasks (2023)

9. Kim, D., Jeong, H., Lee, J., Ghim, J.w., Her, E.S., Lee, S.h., Kim, D., Jeong, H., Lee, J., Ghim, J.w., Her, E.S., Lee, S.h.: Distinct Roles of Parvalbumin- and Somatostatin-Expressing Interneurons in Working Memory Article Distinct Roles of Parvalbumin- and Somatostatin-Expressing Interneurons in Working Memory. Neuron 92(4), 902–915 (2016)

10. Kurikawa, T.: Intermediate Sensitivity of Neural Activities Induces the Optimal Learning Speed in a Multiple-Timescale Neural Activity Model. In: Mantoro, T., Lee, M., Ayu, M.A., Wong, K.W., Hidayanto, A. (ed.) Neural Information Processing. ICONIP 2021, pp. 64–72. Springer, Cham (2021)

11. Kurikawa, T.: Transitions Among Metastable States Underlie Context-Dependent Working Memories in a Multiple Timescale Network. In: Farkaš, I., Masulli, P., Otte, S., Wermter, S. (ed.) Artificial Neural Networks and Machine Learning – ICANN 2021, pp. 604–613 (apr 2021)

12. Kurikawa, T., Kaneko, K.: Multiple-Timescale Neural Networks: Generation of History-Dependent Sequences and Inference Through Autonomous Bifurcations. Frontiers in computational neuroscience 15, 743537 (jun 2021)

13. Maes, A., Barahona, M., Clopath, C.: Learning compositional sequences with multiple time scales through a hierarchical network of spiking neurons. PLoS Computational Biology 17(3) (mar 2021)

14. Manea, A.M.G., Maisson, D.J.N., Voloh, B., Zilverstand, A., Hayden, B., Zimmermann, J.: Neural timescales reflect behavioral demands in freely moving rhesus macaques. Nature Communications 15(1), 2151 (mar 2024)

15. Murray, J.D., Bernacchia, A., Freedman, D.J., Romo, R., Wallis, J.D., Cai, X., Padoa-Schioppa, C., Pasternak, T., Seo, H., Lee, D., Wang, X.J.: A hierarchy of intrinsic timescales across primate cortex. Nature neuroscience 17(12), 1661–1663 (nov 2014)

16. Nishida, S., Tanaka, T., Shibata, T., Ikeda, K., Aso, T., Ogawa, T.: Discharge-Rate Persistence of Baseline Activity During Fixation Reflects Maintenance of Memory-Period Activity in the Macaque Posterior Parietal Cortex. Cerebral Cortex 24(6), 1671–1685 (jun 2014)

17. Pho, G.N., Goard, M.J., Woodson, J., Crawford, B., Sur, M.: Task-dependent representations of stimulus and choice in mouse parietal cortex. Nature Communications 9(1), 2596 (jul 2018)

18. Rikhye, R.V., Gilra, A., Halassa, M.M.: representations enables cognitive flexibility. Nature Neuroscience 21(December) (2018)

19. Runyan, C.A., Piasini, E., Panzeri, S., Harvey, C.D.: Distinct timescales of population coding across cortex. Nature 548(7665), 92–96 (aug 2017)

20. Wasmuht, D.F., Spaak, E., Buschman, T.J., Miller, E.K., Stokes, M.G.: Intrinsic neuronal dynamics predict distinct functional roles during working memory. Nature Communications 9(1), 3499 (dec 2018)

21. Yamashita, Y., Tani, J.: Emergence of functional hierarchy in a multiple timescale neural network model: a humanoid robot experiment. PLoS computational biology 4(11), e1000220 (nov 2008)

